# Proteomic analysis across patient iPSC-based models and human post-mortem hippocampal tissue reveals early cellular dysfunction, progression, and prion-like spread of Alzheimer’s disease pathogenesis

**DOI:** 10.1101/2023.02.10.527926

**Authors:** Yuriy Pomeshchik, Erika Velasquez, Jeovanis Gil, Oxana Klementieva, Ritha Gidlöf, Marie Sydoff, Silvia Bagnoli, Benedetta Nacmias, Sandro Sorbi, Gunilla Westergren-Thorsson, Gunnar K. Gouras, Melinda Rezeli, Laurent Roybon

**Author notes:** **Authors Email address:** Yuriy Pomeshchik, Erika Velásquez, Jeovanis Gil Valdés, Oxana Klementieva, Ritha Gidlöf, Marie Sydoff, Silvia Bagnoli, Benedetta Nacmias, Sandro Sorbi, Gunilla Westergren-Thorsson, Gunnar Gouras, Melinda Rezeli, Laurent Roybon. Department of Neurodegenerative Science, the MiND program, Van Andel Institute, Grand Rapids, MI, USA.

## Abstract

The hippocampus is a primary region affected in Alzheimer’s disease (AD). Because AD postmortem brain tissue is not available prior to symptomatic stage, we lack understanding of early cellular pathogenic mechanisms. To address this issue, we examined the cellular origin and progression of AD pathogenesis in patient-based model systems including iPSC-derived brain cells transplanted into the mouse brain hippocampus. Notably, proteomic analysis of the graft enabled the identification of proteomic alterations in AD patient brain cells, associated with increased levels of β-sheet structures and Aβ42 peptides. Interestingly, the host cells surrounding the AD graft also presented alterations in cellular biological pathways. Furthermore, proteomic analysis across human iPSC-based models and human post-mortem hippocampal tissue projected coherent longitudinal cellular changes indicative of disease progression from early to end stage AD. Our data showcase patient-based models to study the cellular origin, progression, and prion-like spread of AD pathogenesis.

**Highlights:**

- AD patient iPSC-derived brain cells survive in the hippocampus of immunodeficient mice 6 months post-transplantation.
- Proteomic analysis of the grafts reveals profound alterations in cellular biological pathways in iPSC-derived hippocampal cells despite absence of senile plaques.
- Proteomic alterations within transplanted AD iPSC-derived hippocampal cells are reminiscent of early/prodromal AD.
- AD-grafted cells induce proteomic changes in host mouse cells.

## Background

Alzheimer’s disease (AD) is a progressive neurodegenerative disorder of the brain and the most common form of dementia. Pathological hallmarks are amyloid plaques mainly composed of amyloid-β (Aβ) 42 peptides and neurofibrillary tangles consisting of abnormally phosphorylated tau proteins in AD brain tissue [1]. While familial forms of AD have been identified, most cases have an unknown aetiology.

Transgenic animal models overexpressing gene variants associated with AD have uncovered putative mechanisms of cellular pathology [2]. However, these models cannot fully inform on mechanisms underlying early cellular dysfunctions and their progression in patient brain cells, which is needed to successfully develop therapies. Reprogramming of patient-derived somatic cells into induced pluripotent stem cells (iPSCs) [3, 4] has emerged as a powerful tool to model familial and idiopathic AD cellular pathogenesis [5–10]. The cells are human, they carry the genetic makeup of the individual they are generated from, and they are young (embryonic-like). This suggests that cellular phenotypes identified in patient iPSC-derived brain cells could inform about early/prodromal stages of AD.

Numerous studies have demonstrated that iPSC-derived brain cells grown in vitro display molecular and cellular changes associated with AD pathology. We showed that iPSC-derived hippocampal neurons carrying the amyloid precursor protein (APP) p.V717I pathogenic variant, exhibited significant alterations in cellular pathways and networks, coupled to increased intracellular and extracellular Aβ42/40 peptide ratios, synaptic dysfunction, and β-sheet structure formation [7]. A limitation of this approach was that the iPSC-derived brain cells were maintained in an artificial cultured environment, which does not closely mimic the brain environment, despite the cells being grown as spheroids. This important issue was initially addressed by the Goldman group and circumvented by the generation of humanized models, to experimentally study cell autonomous phenotypes within schizophrenia and Huntington’s disease patient glia, in the brain of living mice [11, 12]. A human-rodent chimeric model was recently developed to experimentally study cellular toxicity induced by ApoE4, the strongest genetic risk factor associated with late onset AD, in human cortical neurons in vivo [13]. Additionally, two studies investigated how AD pathology would affect human pluripotent stem cell-derived neurons and astrocytes following transplantation in the AD mouse brain [14, 15]. Lacking however, is a comparison of cell autonomous dysfunctions in AD patient iPSC-derived brain cells grown in vitro and in vivo, and how they differ from those present at end stage of AD.

Here, we generated a chimeric model of early onset/prodromal AD by transplanting AD patient iPSC-derived hippocampal brain cells into the hippocampi of immunodeficient mice. We analyzed cellular pathogenesis in the grafted cells and host tissue 6 months post-engraftment, using advanced imaging, biochemical and liquid chromatography-tandem mass spectrometry (LC-MS/MS) techniques. Finally, we compared the cellular pathways and network dysfunction of the grafted AD cells with those present in AD patient iPSC-derived hippocampal neurons grown in vitro and AD post-mortem hippocampal tissue.

## Methods

All materials were purchased from ThermoFisher Scientific, unless specified.

### Human induced pluripotent stem cell lines

The generation and characterization of human iPSC lines CSC-37N (healthy control) and CSC-17F (APP p.V717I, heterozygous) has been previously reported [7].

### Generation of hippocampal spheroids

After several passages in vitro, iPSCs were differentiated into hippocampal spheroids as previously described [7]. Briefly, to form EBs, iPSC colonies were dissociated using dispase (1 mg/ml) and transferred into ultra-low adherent flasks (Corning) in WiCell medium supplemented with 20 ng/ml FGF2 and 20μM ROCK-Inhibitor Y-27632 (Selleck Chemicals, Munich, Germany). Next day, WiCell was replaced with neural induction medium (NIM) composed of advanced DMEM/F12, 2% B27 Supplement without vitamin A (v/v), 1% N2 Supplement (v/v), 1% NEAA (v/v), 2 mM L-glutamine and 1% Penicillin-Streptomycin (v/v) supplemented with LDN-193189 (Stemgent, 0.1 μM), Cyclopamine (Selleck Chemicals, 1 μM), SB431542 (Sigma-Aldrich, 10 μM) and XAV-939 (Tocris, 5 μM). On the tenth day, the free-floating spheres were transferred to neuronal differentiation medium (NDM) containing Neurobasal^®^ medium, 1% N2 (v/v), 1% NEAA (v/v), L-glutamine and 1% Penicillin-Streptomycin (v/v) supplemented with CHIR-99021 (Stemgent, 0.5 μM) and brain derived neurotrophic factor (BDNF, PeproTech, 20 ng/ml). On the day in vitro 50, spheroids were dissociated into single cells with Trypsin 1X and resuspended in PBS to a final concentration of 100 000 cells/μl for transplantation into the mouse brain.

### In vivo transplantation

For in vivo transplantation, male 6–8 weeks old RAG-2-deficient mice (Janvier Labs, France) were used. The mice were anesthetized with isoflurane (Baxter, Deerfield, IL) in oxygen (initial dose of 5% which was reduced to 1–1.5% for maintenance of surgical depth anesthesia) delivered through a nose mask during the surgery. The body temperature was maintained at 37.0°C during the surgery using a thermostatically controlled rectal probe connected to a homeothermic blanket. The skull was exposed by the skin incision after subcutaneous analgesia with Marcaine (50 μl of 2.5 mg/ml stock solution, Astra Zeneca). Transplantation was stereotaxically performed for each mouse bilaterally through drilled holes in the skull using a 5-μl Hamilton syringe with 32-gauge needle and an injecting minipump (Nanomite Injector Syringe Pump; Harvard Apparatus, Holliston, MA, USA). A volume of 2 μl of cell suspension was injected at a rate of 0.5 μl/min at the following coordinates (from bregma and brain surface): anterior/posterior (AP): −2.0 mm; medial/lateral (M/L): +/−1.5 mm; dorsal/ventral (D/V): −1.7 mm. The needle was left in situ for 7 min after injection before being slowly raised, and the wound was sutured.

### PET imaging

Six months after transplantation, the mice were prepared for imaging in a small animal PET/CT scanner (nanoScan PET/CT, Mediso, Hungary). Eight mice (4 controls and 4 AD) were injected with a mean activity of 14.7 MBq (range: 4.6 – 25.2 MBq) of 18F-flutemetamol intravenously in a tail vein. 70 minutes after injection of the radiopharmaceutical, a PET scan was performed with a scan time of 20 minutes. Prior to and during scanning, the mice were anaesthetized with isoflurane and kept warm via a closed flowing air system in the animal beds. The respiration of the animals was monitored during the whole scan. After the PET scan and sequentially without moving the animal between scans, a 15-minute CT scan was performed at 45kV and a 900 ms exposure time. PET image reconstruction was performed using the Tera-Tomo™ three-dimensional (3D) PET image reconstruction engine with 4 iterations and 6 subsets and a voxel size of 0.4×0.4×0.4 mm3 (Mediso, Hungary). The CT image was used for attenuation correction and as anatomical reference during analysis. One of the control animals moved during the PET scan, and thus had to be excluded from the analysis. Image analysis was performed using the software Vivoquant™ 4.0patch1 (InviCRO, Needham, MA, USA). Region of interests (ROIs) were drawn in the fused PET/CT images and values of activity uptake and volume were extracted for calculation of %IA/g for each animal.

### Rodent brain tissue collection and processing

Six months after transplantation, the mice were transcardially perfused with PBS. The brain tissues containing grafts from the left and right brain hemisphere were dissected, cut into two halves each, snap frozen on liquid nitrogen and stored at −80°C for further biochemical analysis. For immunohistochemistry, the mice were transcardially perfused with PBS followed by perfusion with 50 ml of 4 % paraformaldehyde (PFA). The dissected brains were post-fixed in the same fixative overnight at 4°C. After fixation, brains were cryoprotected in 30% sucrose (Sigma-Aldrich) for 48 hours at 4°C before being cut into 30 μm thick coronal serial sections on a sliding microtome (Leica Biosystems, Wetzlar Germany). The free-floating brain sections were then stored in antifreeze solution at −20°C until immunohistochemical staining.

### Human postmortem brain tissue processing

The human postmortem brain samples, hippocampi from AD patients and non-demented controls (Supplementary table 6) were obtained from The Netherlands Brain Bank, Netherlands Institute for Neuroscience, Amsterdam (open access: www.brainbank.nl). All Material has been collected from donors for or from whom a written informed consent for a brain autopsy and the use of the material and clinical information for research purposes has been obtained by the NBB. The frozen hippocampi samples were cut into 20 μm thick sections on a cryostat (Leica Biosystems) and then stored at −80°C until immunohistochemical staining or biochemical analysis.

### Immunostainings, microscopy, and image analyses

For immunohistochemistry, frozen mouse or human sections were air-dried for 1 hour, washed 3 times in PBS, blocked for 1hr at RT with 5% normal donkey serum (NDS, VWR) in PBS with 0.25% Triton-X (Sigma-Aldrich) and incubated overnight with target primary antibodies (Supplementary table 7) prepared in 5% NDS in PBS with 0.25% Triton-X at 4°C. Prior to blocking, the human sections were fixed with 4% PFA for 1hour at RT. On the next day, the sections were incubated with appropriate Alexa-fluor 488 or 555-conjugated secondary antibodies (Supplementary table 7) in PBS for 2hr at RT in the dark. Cell nuclei were counterstained with DAPI (1:10,000). Congo Red staining was performed using Congo Red Staining Kit (Merck) following the manufacturer’s instructions. Image acquisition was performed using inverted epifluorescence microscope LRI-Olympus IX-73 equipped with fluorescence filters. Automated quantitative image analysis was performed with the ImageJ software or MetaMorph Software V7.6 (Molecular Devices) using the Multi-Wavelength Cell Scoring application. For a specific marker, positive cells were identified as having signal intensity above the selected intensity threshold. Intensity thresholds were set blinded to sample identity.

### MSD MULTI-ARRAY Assay

Aβ40 and Aβ42 levels were quantified in brain lysates using the multiplex Aβ Peptide Panel 1 kit (6E10, MesoScale Discovery, USA) according to the manufacturer’s protocol.

### Western blot analysis

For WB, samples were mixed with loading buffer, heated at 70 °C for 10 min and loaded into Bolt 4–12% Bis-Tris Plus gels. After electrophoresis the samples were transferred to polyvinylidine difluoride (PVDF) membranes. Membranes were blocked in PBS containing 0.1% Tween-20 (PBST) and 5% milk or 5 % BSA and incubated with target primary antibodies (Supplementary table 8), overnight. Next day, membranes were incubated with HRP-conjugated secondary antibodies (Supplementary Table 8) for 1 h diluted in PBST and 5% milk or 5% BSA. The blots were then visualized using Pierce Enhanced Chemo-Luminescence solution and imaged in a Biorad Chemi-Doc chemo-luminescence system (BioRad).

### Dot blot analysis

For Dot blot analysis, 20 ug of proteins of each sample was deposited directly onto PVDF membranes. The blots were incubated with OC antibody (Sigma-Aldrich, Sweden, #AB 2286, 1:5000) that targets amyloid fibrils (Kayed et al., 2007), overnight. The next day, the membranes were washed and incubated with an anti-mouse HRP-conjugated secondary antibody (R&D Systems; 1:1000) for 1 h. The blots were then visualized using Pierce Enhanced Chemo-Luminescence solution and imaged in a Biorad Chemi-Doc chemoluminescence system (BioRad).

### Fourier transformed infrared micro-spectroscopy

For FTIR micro-spectroscopy analyses, fresh frozen mouse brain tissues containing human grafts were cut into 16 μm thick sections on a cryostat. Sections were mounted on the 1 mm thick CaF2 round 10 mm spectrophotometric windows. Infrared spectra were taken from RANDOM areas of the section at the SMIS beamline of the SOLEIL synchrotron (SMIS beamline, France) using a Thermo Fisher Scientific Continuum XL FTIR microscope through a 32× magnification, 0.65 NA Schwarzschild objective. For the collection, parameters were at spectral range 1000-4000 cm^−1^, in transmission mode at 4 cm^−1^ spectral resolution, with 10 μm × 10 μm aperture dimensions, using 256 codded scans. Background spectra were collected from a clean area of the same CaF2 window. All measurements were made at room temperature. For analysis of FTIR spectra OPUS software (Bruker) and Orange (University of Ljubljana) were used and included atmospheric compensation. Derivation of the spectra to the second order using Savitsky-Golay of 3^rd^ polynomial order 3 with 9 smoothing points, was used to unmask the number of discriminative features and to eliminate a contribution of a baseline.

### Protein extraction and determination

Brain tissues for immunoassays were resuspended in 200 μl of ice-cooled lysis buffer (50 mM Tris-buffer, 150 mM NaCl, 0.05% Tween-20, protease, and phosphatase inhibitor cocktail (1X)). Then, the proteins were extracted by sonication executing 40 cycles of 15s on and 15 s off at 4 °C using a Bioruptor plus (model UCD-300, Diagenode). After lysis, samples were centrifuged at 4 °C, 20.000 g for 20 minutes, supernatants were collected and stored at −80 °C until further use. Total protein amount was determined using the Pierce™ BCA Protein Assay Kit according to the manufacturer instructions. Protein extraction of the hippocampal spheroids for proteomic analysis was performed as previously described (Pomeshchik Y et al., 2020). Proteins from the hippocampal graft and hippocampal post-mortem brain tissues were extracted using a lysis buffer of 25mM DTT, 10 w/v% SDS in 100mM Triethylammonium bicarbonate (TEAB). Samples were sonicated using 40 cycles of 15s on/off at 4 °C in the Bioruptor plus (model UCD-300, Diagenode) after boiling at 99 °C for 5 minutes. Samples were centrifuged at 20000 g for 15 min at 18°C, and the supernatant was collected. Protein concentrations were measured using the Pierce™ 660nm Protein Assay with ionic detergent compatibility reagent (Thermo) and preserved at −80 °C until further use.

### Protein digestion for proteomic analysis

Samples were alkylated with 50 mM IAA for 30 min in the dark at room temperature. Protein digestion was performed using the S-Trap™ 96-well plate following the manufacturer’ instructions (PROTIFI. S-Trap™ 96-well plate digestion protocol. https://files.protifi.com/protocols/s-trap-96-well-plate-long-1-4.pdf.) Briefly, 95μL of 50 mM TEAB containing LysC (enzyme: substrate,1:50) was added to each sample and incubated for 2 h at 37 °C, followed by the addition of 30 μL of 50 mM TEAB containing Trypsin (enzyme: substrate,1:50). Samples were incubated overnight, at 37 °C. The reaction was stopped by acidifying the samples with 40 μL of formic acid (FA). Peptides were dried in a speed-vac and resuspended in 0.1% trifluoroacetic acid (TFA)/2%acetonitrile (ACN). Peptide concentrations were measured using the Pierce™ Quantitative Colorimetric Peptide Assay.

### LC-MS/MS analysis and database searching

Two micrograms of peptides were separated on an Ultimate 3000 RSCLnano pump system using an Acclaim PepMap100 C18 (5 μm, 100 Å, 75 μm i.d. × 2 cm, nanoViper) trap column and an EASY-spray RSLC C18 (2 μm, 100 Å, 75 μm i.d. × 50 cm) analytical column coupled to a Q Exactive HF-X mass spectrometer. The flow rate was set to 300 nl/min for 120 min, with a column temperature of 60°C. For the chromatographic gradient 0.1% FA as solvent A and 80% ACN/ 0.1%FA as solvent B were used. Mass spectra were acquired using the data-dependent acquisition mode. Full scans were collected at 120,000 resolution with a maximum injection time (IT) of 100 ms and a target AGC value of 3e06. The 20 most intense ions were selected for fragmentation using a normalized collision energy (NCE) of 28. MS2 scans were acquired with a resolution of 15,000, a target AGC value of 1e05, and a maximum IT of 50 ms. The data were searched against the UniProt human database using the SEQUEST HT algorithm in the Proteome Discoverer (PD) 2.3 software. The hippocampal graft samples were additionally searched against the *Mus musculus* database (UniProt). Cysteine carbamidomethylation was considered as static modification, methionine oxidation and the N-terminal protein acetylation were included as dynamic modifications. A maximum of 2 missing cleavages was allowed. The precursor mass tolerance was set to 10 ppm, and the fragment mass tolerance to 0.02 Da. A correction of 1% false discovery rate (FDR) at both peptide and protein levels was applied.

### Statistical and biological pathway analyses

Protein intensities were normalized by log2 transformation and the subtraction of median intensities using the software Perseus 1.6.5.0. Principal component analysis of the global proteome was also performed in Perseus 1.6.5.0. To determine statistically significant differences between conditions a two-tailed t-test (p-value:0.05) with Benjamini-Hochberg (FDR: 0.05) correction for multiple testing was applied. A cutoff of + 0.5 was also set for the log2 fold-change (AD/Control) in the post-mortem brain tissue samples. To select proteins significantly altered across the different AD models a one-way ANOVA (p-value:0.05) with Benjamini-Hochberg (FDR: 0.05) correction was applied together with the log2 fold-change (AD/Control) cutoff. The Tukey’s honest significant difference test was used as a post-hoc test (FDR< 0.05). Additionally, hierarchical clustering analysis was performed using the Pearson correlation distance. For the 1D annotation enrichment analysis of the quantified proteins the Benjamini-Hochberg FDR value of 0.02 was set as a significance limit. The detection of relevant biological pathways was performed in the Metascape [16] incorporating the Gene Ontology Consortium, Reactome Gene Sets, KEGG Pathway, and PANTHER Pathway databases. Enrichment processing included the gene prioritization by evidence counting strategy with a p-value cutoff of 0.01.

Finally, data derived from the immunoassays were analyzed using GraphPad Prism 7 software and presented as mean ± S.E.M. Unpaired two-tailed t-test was used to compare the two groups. A p-value of < 0.05 was considered significant.

## Results

### Human iPSC-derived hippocampal brain cells survive in the mouse brain and express neuronal and astrocytic markers

To investigate the effect of an APP pathogenic variant in human hippocampal brain cells *in vivo*, we differentiated iPSCs lines generated from a non-demented female individual and an AD female patient carrying the most common missense variation of the APP gene (APP p.V717I), into hippocampal spheroids [7]. Fifty-day old spheroids containing hippocampal neurons and neural progenitors were dissociated into single cells and transplanted into the hippocampi of 3-month-old Rag2 immunodeficient mice. Immunohistochemical analysis of the graft 6-month post-transplantation revealed that the human cells had survived and integrated in the host hippocampi (Figure 1b-d). We estimated that 5% of all human cells were still actively dividing (Figure 1d and Figure S1). Amongst the human grafted cells, 40%-65% and 30%-40% stained positive for MAP2 and Doublecortin, respectively (Figure 1e and f). While most human neurons expressed T-Box Brain Transcription Factor 1 (TBR1) and Zinc Finger and BTB Domain Containing 20 (ZBTB20) markers of hippocampal identity, few were positive for PROX1 (Figure 1e and f). Immunohistochemistry for STEM123 and ionized calcium binding adaptor molecule 1 (IBA1) revealed the presence of human astrocytes and the absence of human microglia, respectively (Figure 1f and Figure S1). Interestingly, the number of host astrocytes was greater in the AD hippocampal grafts (ADHG), compared to control hippocampal grafts (CHG). It is worth mentioning that as opposed to mouse astrocytes, the number of mouse microglial inside the graft did not differ between AD and control grafts (Figure 1f and Figure S1).

**Figure 1.**
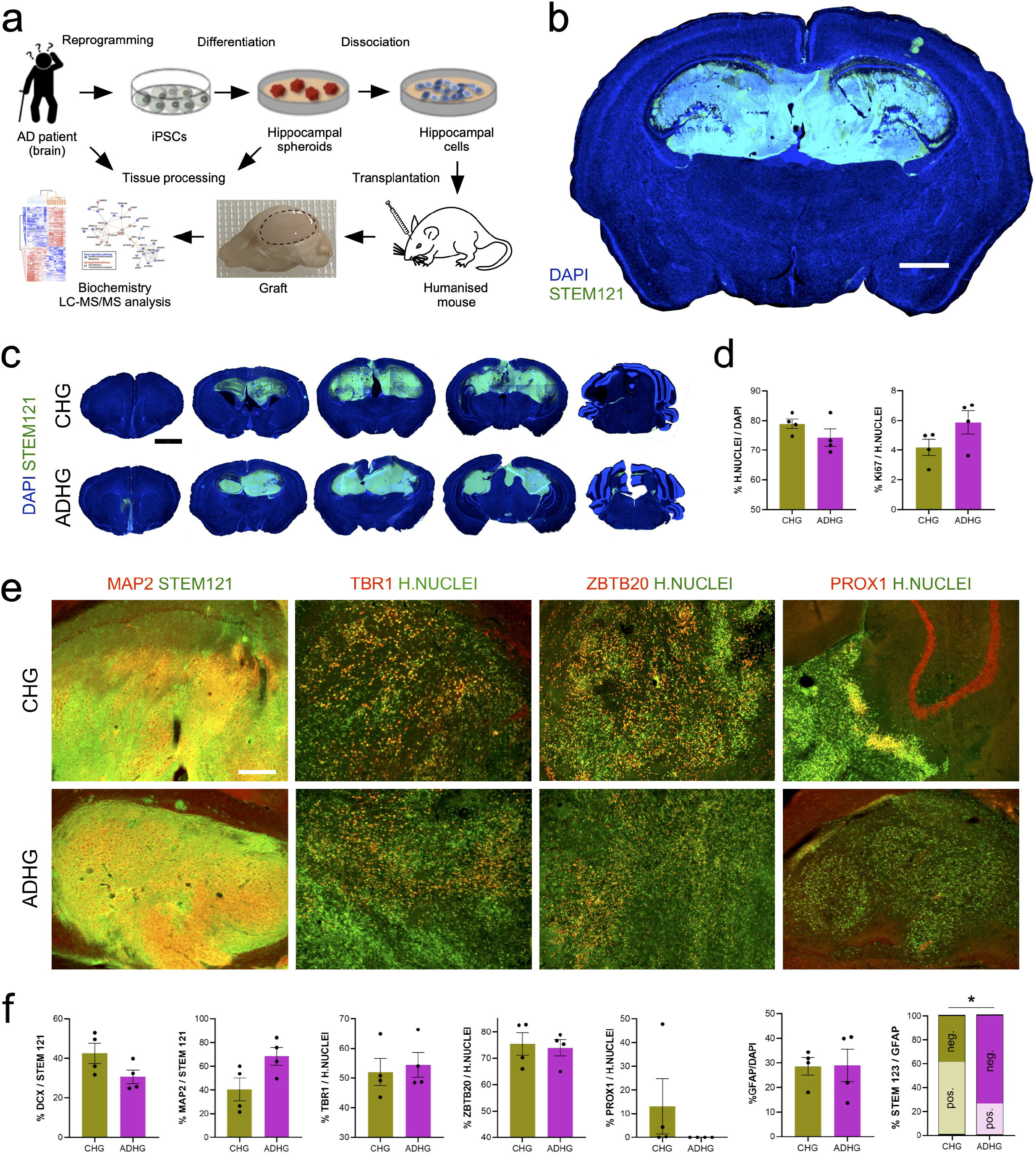
Characterization of the iPSC-derived human grafts (HG) in the mouse brain. (a) Schematic representation of the experimental workflow. (b and c) Immunostaining for human cytoplasmic marker STEM121 and nuclear marker DAPI in control and AD grafts six months after transplantation into mouse hippocampus. Scale bars: 1 mm (b) and 2 mm (c). ADHG: AD human graft; CHG: Control human graft. (d) Quantification of human Nuclei marker-positive cells expressed relative to the total number of DAPI-labeled cells and Ki-67-positive cells expressed relative to the total number of human Nuclei marker-positive cells in control and AD iPSC-derived human grafts. Results are presented as mean ± S.E.M. N = 4 animals. Statistical analysis by two-tailed t-test. (e) Immunostaining for human Nuclei marker and neuronal markers MAP2, TBR1, and hippocampal markers ZBTB20 and PROX1 in control and AD iPSC-derived human grafts. Scale bars: 200 μm. (f) Quantification of DCX-, MAP2-, TBR1-, ZBTB20-, PROX1-, GFAP- and STEM123-positive cells in control and AD iPSC-derived human grafts. Results are presented as mean ± S.E.M. N = 4 animals. P value: *p < 0.05. Statistical analysis by two-tailed t-test.

Together, these data show that the grafts were composed of neuroblasts, neurons and astrocytes, and that neurogenesis was still ongoing 6-month post-transplantation.

### AD cellular pathology identified in 6-month-old graft despite absence of senile plaques

The formation of senile plaques caused by the accumulation and aggregation of mainly Aβ42 peptides forming amyloids in the brain extracellular space, is one of the main hallmarks of AD pathology [17]. Positron emission tomography (PET)-mediated imaging of Aβ fibrils in the brain using tracer flutemetamol (^18^F) is very efficient at detecting amyloid deposits in the brain of AD or suspected AD patients[18]. The technique has also been used to define when, during the course of the disease, mouse models of AD develop Aβ plaques [19]. To detect whether Aβ deposition had occurred in 6-month-old ADHG, we subjected the humanized mice to ^18^F-PET imaging. No difference in signal activity was observed in the brain of the animals transplanted with AD hippocampal brain cells compared to those transplanted with control hippocampal brain cells (Figures 2a and b). This data, which suggested the absence of Aβ plaques in the transplanted animals, prompted us to examine whether amyloid fibrils could be identified in the AD grafts using alternative techniques. In corroboration with the ^18^F-PET imaging data, no staining was identified in either the AD or control graft, after processing of the tissue with either Aβ antibody H31L21 immunohistochemistry or Congo-red staining, compared to AD mouse brain tissue used as positive control (Figures 2c and d).

**Figure 2.**
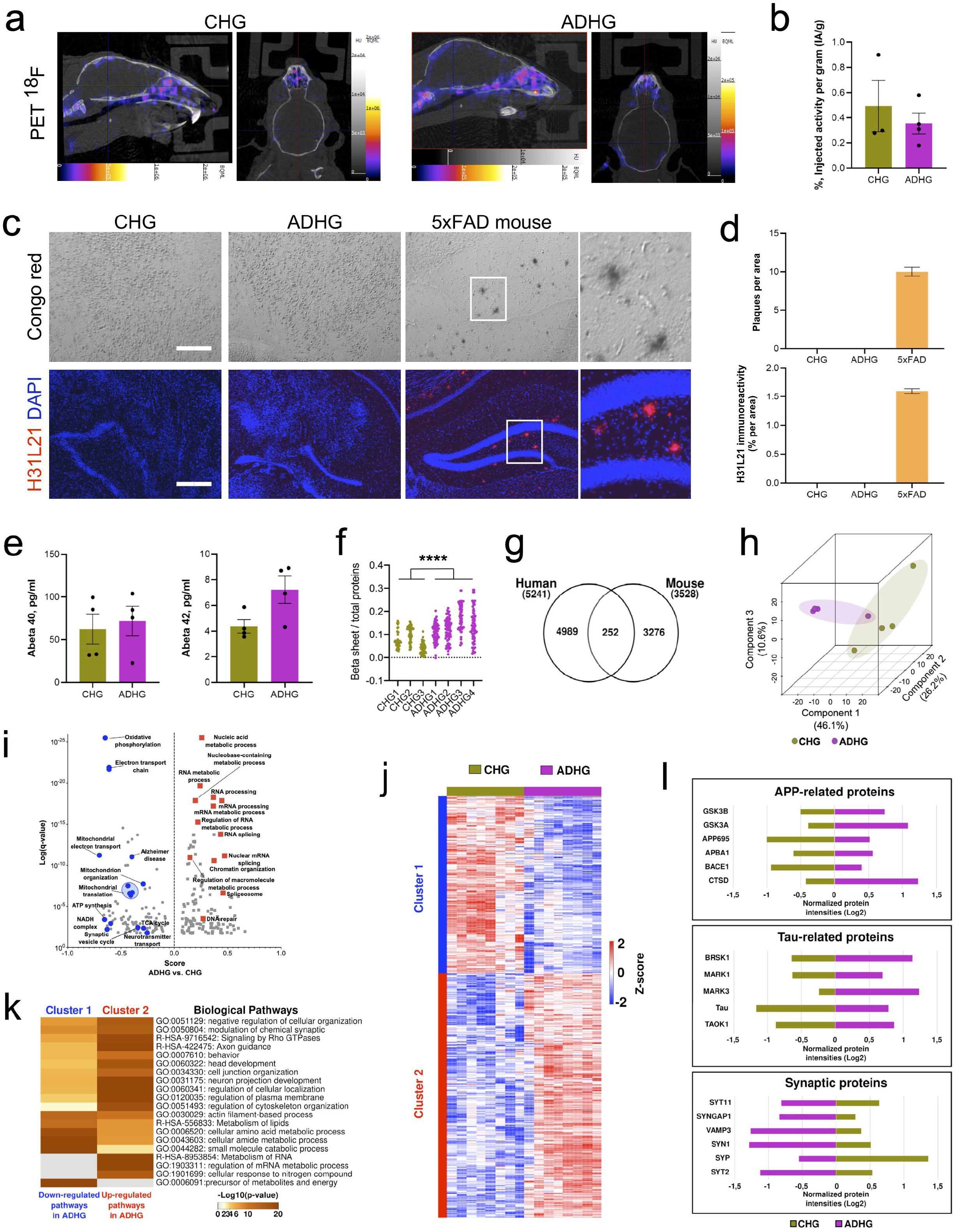
AD human grafts exhibit significant proteomic alterations, but do not show main AD hallmarks. (a and b) Overview of representative PET/CT images of ^18^F-flutemetamol in control and AD iPSC-derived grafts six months after transplantation into mouse hippocampus. Sagittal and axial views showed no specific uptake related to AD in the brain. Color scale bar shows (from black to white) activity values in PET ^18^F-flutemetamol image. The diagram shows relative uptake values presented as %IA/g. Results are presented as mean ± S.E.M. N = 3-4 animals. (c and d) Characterization of Aβ deposition in hippocampi of 6-month-old 5xFAD transgenic mice and control and AD iPSC-derived grafts. Congo Red staining and immunostaining for H31L21 show presence of Aβ deposits only in 5xFAD mice. Scale bars: 200 μm. Results are presented as mean ± S.E.M. N = 4 animals. (e) Accumulation of Aβ measured by MSD in control and AD iPSC-derived human grafts. Results are presented as mean ± S.E.M. N = 4 animals. Statistical analysis by two-tailed t-test. (f) Diagram reflecting the β-sheet structure content in control and AD iPSC-derived human grafts as shown by the absorbance ratios 1,628 cm^−1^, a band characteristic for β-sheet structure, to 1,656 cm^−1^, a band typically assigned to α-helical or random structures, main components of protein structures used here as a normalization parameter. Results are presented as individual values. n = 42 - 81 spectra per sample for N = 3-4 animals. P value: **** = P<0.0001. Statistical analysis by two-tailed t-test. (g) Venn diagram representing the number of identified proteins exclusively derived from AD iPSC-derived human grafts and/or from host mouse brain tissue. N = 4 animals. (h) Principal component analysis using the normalized intensities of proteins quantified in control and AD iPSC-derived human grafts. (i) 1D annotation enrichment analysis of quantified proteins in control and AD iPSC-derived human grafts (Benjamini-Hochberg; FDR: 0.02). Underrepresented pathways in ADHG (blue) are mainly enriched by mitochondrial function. Overrepresented biological pathways in ADHG (red) are mostly associated with the dysregulation of RNA metabolism. (j) Hierarchical clustering analysis of dysregulated proteins in AD iPSC-derived human grafts (Two-tailed t-test; p-value < 0.05; Benjamini-Hochberg FDR: 0.05) using Pearson correlation distance. The abundance of protein groups decreased and increased is represented in blue and red, respectively. (k) Biological pathways enriched in the protein clusters of the hierarchical clustering analysis of AD iPSC-derived human graft proteome (p-value < 0.01). P-values are denoted as the – log10 (p-value). The grey color symbolizes the absence of that specific biological pathway. l) Cluster bar representation of significantly altered proteins linked to well-established markers of early molecular dysfunction in AD (Two-tailed t-test; p-value < 0.05; Benjamini-Hochberg FDR: 0.05).

Because the human cells that compose the graft are young, it is possible that they exhibit signs of AD cellular pathology prior to senile plaque formation. We next utilized MSD multiarray to measure the amount of Aβ40 and Aβ42 peptides present in the human grafts. Levels of Aβ40 peptide were not different between CHG and ADHG. However, levels of Aβ42 showed a trend towards significantly increased levels in ADHG compared to CHG (Figure 2e). Additionally, we found increased levels of β-sheet structures measured by Fourier transform infrared (FTIR) micro-spectroscopy. Increased levels of β-sheet structures preceding the formation of amyloid aggregates was previously identified in AD patient iPSC-derived hippocampal spheroids in vitro and in young mouse models of AD [7, 20].These data which suggested that cellular pathology might be initiating, prompted us to examine if and what cellular pathways and networks were changed in the AD patient brain cells. Towards this end, we assessed the proteomic landscape in the grafts, by taking advantage of quantitative protein analysis by LC-MS/MS (Table S1). Using the *Homo sapiens* and *Mus Musculus* databases (Uniprot) integrated to the pipeline of mass spectra searching, we were able to dissect the proteome map derived from the human and mouse cells that composed the grafts. The quantification of unique peptides (at least two per protein) successfully discriminated the human graft protein profile from the mouse brain proteome, with minimal overlap in protein quantification (Figure 2g). Principal component analysis (PCA) based on the quantified human global proteome clearly discriminated ADHG from CHG groups (Figure 2h). Enrichment analysis showed over-representation of RNA metabolic pathways in human cells that composed the ADHG, while biological pathways related to mitochondrial and synaptic function were under-represented (Figure 2i).

Further analysis allowed us to scrutinize which proteins had altered levels in human cells composing the ADHG compared to CHG (Table S2). Hierarchical clustering analysis allowed for the identification of two main protein clusters (Figure 2j). Cluster 1 encompassed proteins whose level was decreased in the human cells composing the ADHG, featured by alterations in energy-related metabolic pathways, and lipid and amino acid metabolism. Increased proteins grouped in the second cluster, characterized by the increment of RNA metabolism, plasma membrane and cytoskeleton regulation, membrane trafficking, and Rho GTPases signaling (Figure 2k). Since altered APP processing, increased Tau phosphorylation and impaired synaptic transmission play pivotal roles in AD [21], we next focused on several proteins involved in these processes. In addition to observe higher levels of APP and Tau, we found increased levels of proteins involved in APP processing and Tau phosphorylation, coupled to a decrease of several synaptic proteins in human cells composing the ADHG, compared to CHG (Figure 2l).

Taken together, these data suggest that important pathways and networks are altered in human cells in the ADHG; and that they coincide in time with the formation of β-sheet structures and accumulation of APP and Tau but precede the formation of senile Aβ plaques.

### Comparison of proteomic changes across human AD hippocampal spheroids, human cells in ADHG and AD patient post-mortem hippocampal tissue suggests a transition of cellular changes from early to end-stage AD cellular pathology

To better understand which disease stage was modelled within the human grafted AD cells, we compared the proteomic changes taking place in human cells in ADHG with those in APP variant hippocampal spheroids (ADHS) aged for 100 days in vitro [7] and human AD hippocampal postmortem brain tissue (ADHPMBT; Table S3), representing early and end stage AD, respectively. First, we confirmed the presence of amyloid plaques in ADHPMBT sections by immunohistochemistry and dot-blot assay, the latter using OC antibody that specifically recognizes amyloid fibrils [22] (Figures 3a-c). Additionally, immunohistochemistry and Western blotting performed with AT8 antibody showed increased phosphorylation of Tau in ADHPMBT compared to control tissue obtained from non-demented healthy controls (Figures 3a, b and d).

**Figure 3.**
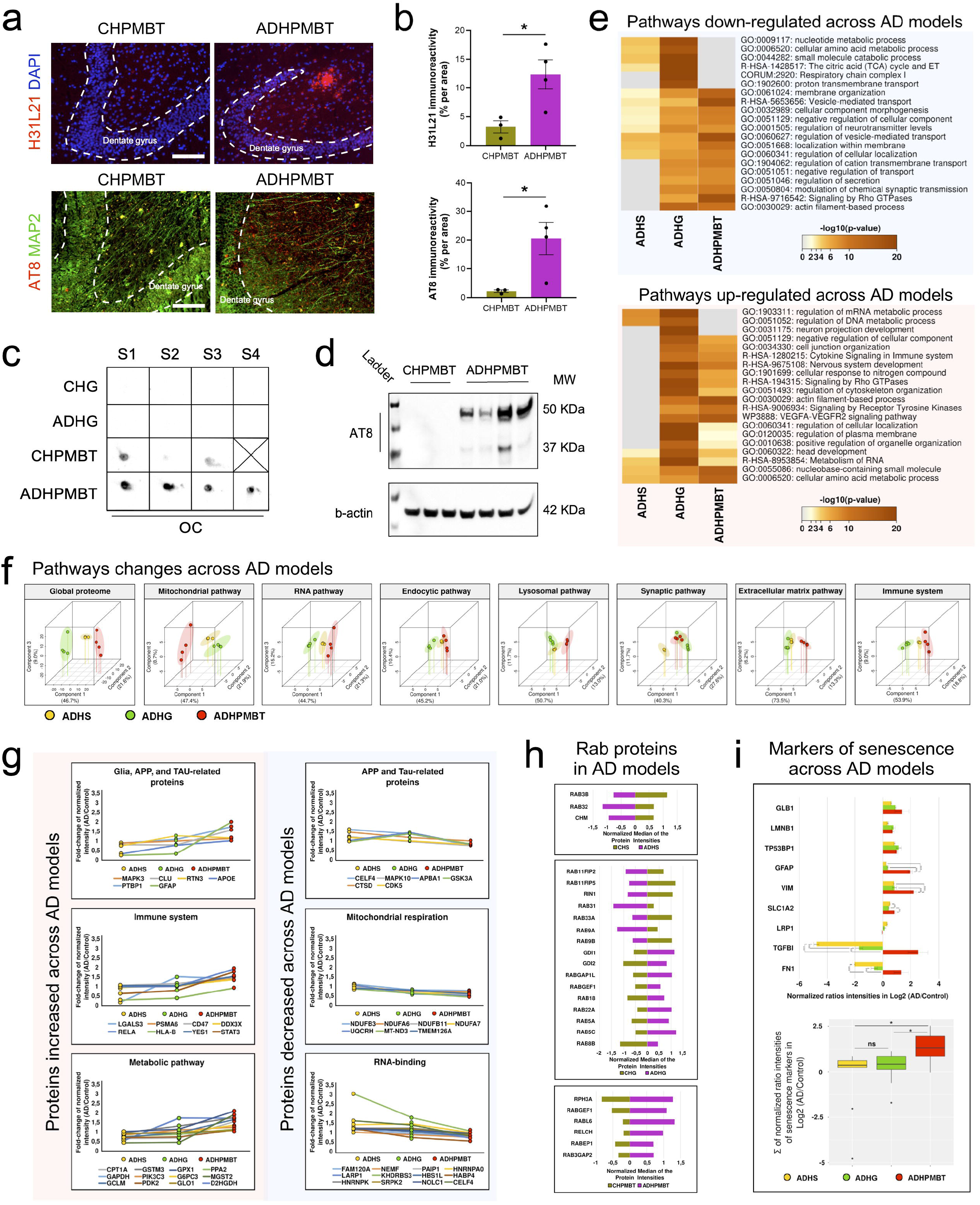
Label-free quantitative proteomics reveals common and distinctive alterations in iPSC-based AD models and postmortem hippocampi from AD patients. (a and b) Characterization of amyloid-β deposition and phosphorylation of tau protein in postmortem hippocampi of AD patients and non-demented controls. Immunostaining for H31L21 show presence of Aβ deposits in postmortem hippocampi of AD patients. H31L21 immuno-positive area was quantified relative to the total area. Immunostaining for AT8 show increased tau phosphorylation in postmortem hippocampi of AD patients. AT8 immuno-positive area was quantified relative to the total area. Results are presented as mean ± S.E.M. N = 3-4 animals. P value: *p < 0.05. Statistical analysis by two-tailed t-test. CHPMBT: control human postmortem brain tissue; ADHPMBT: AD human postmortem brain tissue. (c) Dot blot immunoassay performed using OC antibody specific to amyloid fibrils showing their presence in postmortem hippocampi of AD patients and non-demented controls, but not in human grafts (b). Scale bars: 200 μm. (d) Western blotting analysis of phosphorylation of tau protein with actin blot included as a loading control. N = 3-4 animals. (e) Comparison of the biological pathways dysregulated in AD iPSC-derived human spheroids (ADHS), AD iPSC-derived human grafts, and AD human postmortem brain tissues with respect to the control conditions (p-value < 0.01). P-values are denoted as the −log10 (p-value). The grey color symbolizes the absence of that specific biological pathway. (f) Principal component analysis of the normalized protein intensities that participate in key signaling pathways affected in AD. The proteomic profiles of each biological pathway are unique for each AD iPSC-derived model and PMBT, showing a clear separation between them. (g) Selected proteins significantly altered across the different AD models. Median fold-change (AD/Control) of normalized intensities; One-way ANOVA p-value < 0.05; Benjamini-Hochberg FDR < 0.05); Tukey HSD FDR< 0.05. (h) Quantification of dysregulated Rab proteins in each AD model. Bar graphs show the median of normalized protein intensities. Statistical significance was assessed by a two-tailed t-test; p-value < 0.05; Benjamini-Hochberg FDR < 0.05 and the log2 fold-change cutoffs of + 0.5. (i) Quantification of dysregulated markers of senescence across the different AD models. Oneway ANOVA p-value < 0.05; Benjamini-Hochberg FDR < 0.05; Tukey HSD FDR< 0.05.

Next, we reasoned that the presence of amyloid deposition and hyperphosphorylated Tau in ADHPMBT seen in end stage AD, should be paralleled by proteomic alterations in line but more advanced than those identified in the ADHG and ADHS. To validate this hypothesis, we mapped the proteome in the two iPSC-based models and ADHPMBT and examined the main commonly dysregulated protein pathways (Figure 3e). Pathways linked to DNA and mRNA metabolism were increased in the two iPSC-based models of AD, but not in ADHPMBT. In contrast, cytokine signaling, membrane trafficking, and cytoskeleton organization were upregulated in the human cells composing the ADHG and ADHPMBT, but not in the ADHS. Mitochondrial metabolism imbalance was shared between ADHS and human cells composing the ADHG, while decrease in transport regulation and neurotransmission was shared between human cells composing the ADHG and ADHPMBT (Figure 3e).

The acquisition of the protein profiles in the iPSC-based models and ADHPMBT allowed us to further interrogate if they shared proteomic alterations. The common total proteins quantified in iPSC-based models and ADHPMBT depicted distinctive and unique signatures of dysregulated biological pathways. These were not only identifiable at the global level, but also when analysing each pathway separately (Figure 3f). To compare the relative abundance of AD-related proteins across the three AD systems, we expressed their amount as a fold change AD/control (Table S4). The fold change of glial fibrillary acidic protein (GFAP), an astrocytic cytoskeletal protein, known to be markedly upregulated in brains of AD patients [23], increased from ADHS and ADHG to ADHPMBT. In line with this, the fold change of signal transducer and activator of transcription 3 (STAT3), a transcription factor regulating many pathways associated with astrogliosis [24, 25] was elevated from ADHS to ADHPMBT. Similarly, the fold change of several proteins involved in APP processing, Tau phosphorylation, immune system activation, and metabolic-related proteins was minimal in ADHS and maximal in ADHPMBT, reflecting progression of AD cellular pathology from AD iPSC-based models to postmortem ADHPMBT (Figure 3g). In contrast, the fold change of proteins involved in mitochondrial respiration and RNA-binding, as well as other proteins related to APP and Tau decreased from ADHS to ADHPMBT. These data likely reflected events that occur early in AD cellular pathogenesis (Figure 3g).

Recently, human iPSC-derived neurons whose genome was edited to express different genes leading to familial AD, revealed endosomal dysfunction as a shared disease-associated cellular phenotype [26]. In line with this work, we examined the levels of several members of the Rab superfamily, small GTPase proteins containing many key regulators of endosome trafficking and remodelling [27], in all three AD systems. While only three Rab proteins were dysregulated in ADHS, 16 members of the Rab superfamily were altered in ADHG and 6 in ADHPMBT. Among them, RABGEF1, a guanine nucleotide exchange factor for RAB5 [28], a key regulator of endosome fusion and trafficking [26, 29], was elevated in both ADHG and ADHPMBT. Interestingly, two isoforms of RAB5, RAB5A and RAB5C were increased only in ADHG. Finally, to evaluate the advance of cellular pathogenesis in the graft, we analyzed the abundance of several senescence markers across all three AD systems. Because proteomics of the samples was not performed at the same time, we expressed their amount as a fold change (AD/control). Data analysis showed prevalence of senescence-related proteins in ADHPMBT compared to ADHG and ADHS, and their mild increase in ADHG compared to ADHS (Figure 3i).

Because significant protein alterations are shared between human cells in the ADHG and ADHPMBT, and senile plaques are absent in ADHG, our data suggests that the cellular pathology within human cells in the ADHG may be reminiscent of the early/prodromal stage.

### Dysfunction of cellular biological pathways identified in host brain cells

Like many neurodegenerative diseases, AD is considered a prion-like disease [30–32]. To address the issue, we asked whether the grafted human APP mutant cells could potentiate cellular pathways and network alterations in surrounding mouse cells. Proteomic analysis allowed us to discriminate 3276 mouse host proteins (Figure 2g). The PCA analysis clearly showed separation of the proteome in the mouse tissue adjacent to the AD human graft (ADHG-MT), compared to control (CHG-MT) (Figure 4b; Table S5). Clustering analysis identified two main groups of dysregulated pathways in the ADHG-MT (Figure 4d). Cluster 1 exhibited upregulated pathways involved in regulating mRNA stability, transmembrane regulation, and synapse organization, whereas cluster 2 showed downregulated pathways mainly linked to metabolism, such as the citric acid cycle and electron transport chain (Figure 4d).

**Figure 4.**
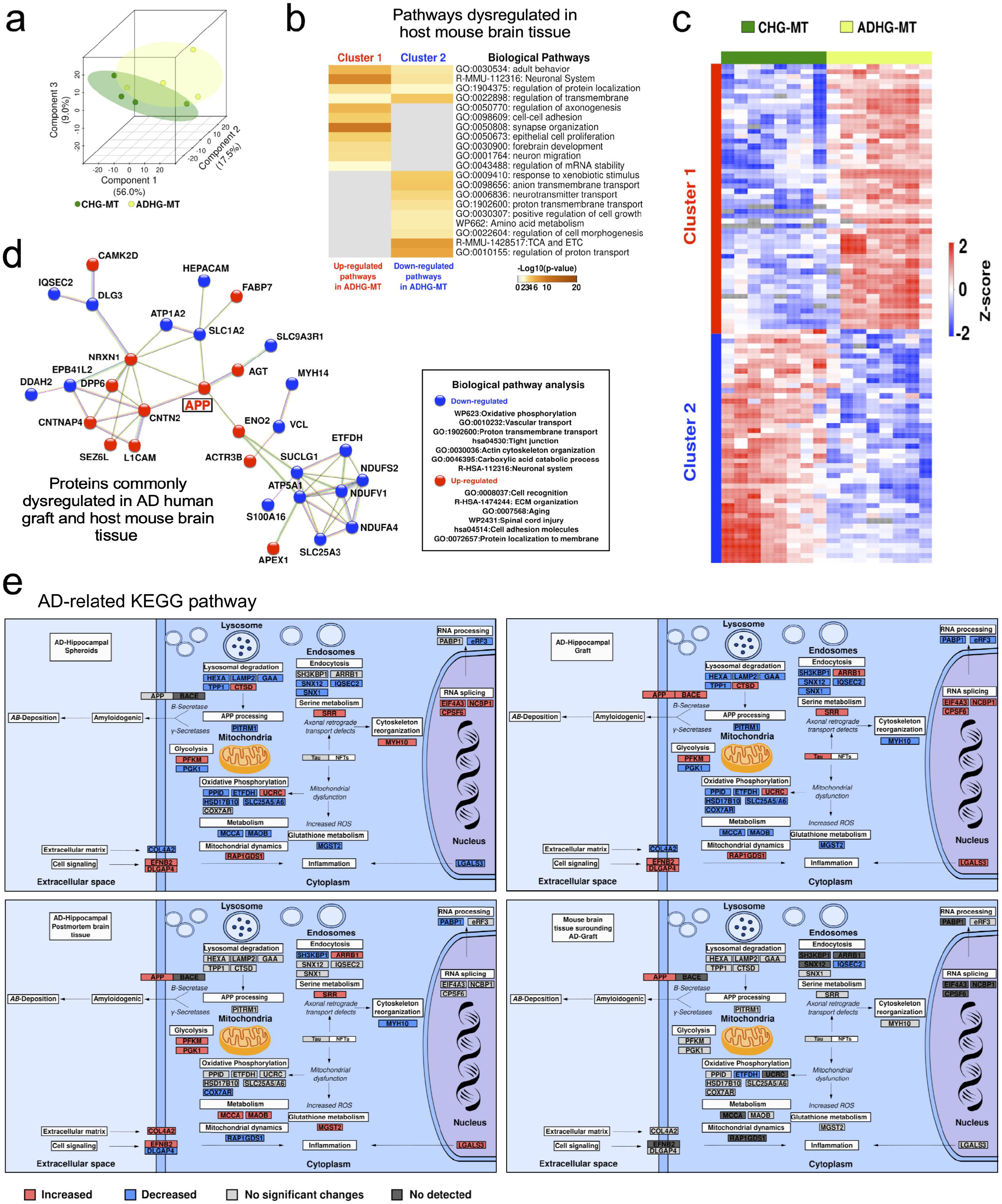
Label-free quantitative proteomics reveals alterations in the host mouse brain tissue induced by AD human grafts. (a) Principal component analysis of murine proteins identified in control and AD-grafted mouse hippocampi. CHG-MT (control human graft-mouse tissue); ADHG-MT (AD human graft-mouse tissue). (b) Hierarchical clustering analysis of dysregulated host mouse proteins (Two-tailed t-test; p-value < 0.05; Benjamini-Hochberg FDR < 0.05) using Pearson correlation distance. The abundance of protein groups decreased and increased is represented in blue and red, respectively. (c) Pathway analysis of the protein clusters in C (p-value < 0.01). P-values are denoted as the −log10 (p-value). The grey color symbolizes the absence of that specific biological pathway. (d) Interaction network and pathway enrichment of the commonly dysregulated proteins between AD iPSC-derived human grafts and the host mouse tissues (Two-tailed t-test; p-value < 0.05; Benjamini-Hochberg FDR < 0.05). Proteins with decreased and increased abundances are represented in blue and red, respectively. (f) Mapping of the dysregulated proteins participating in the AD pathway reported in the KEGG database. The analysis was centered on the dysregulated proteins (AD vs. Control) in the ADHG model in comparison to ADHS, ADHPMBT, and ADHG-MT. Protein groups with decreased and increased abundances are represented in blue and red, respectively. Proteins with no significant changes in their abundance and not detected are represented in grayscale.

We next examined how proteins whose levels were changed in both human and mouse cells connected with each other. We used the UniProt tool to convert the *Mus musculus* entries to *Homo sapiens*. The data analysis demonstrated an overlap in crucial AD-related proteins and pathways (Figure 4e). Notably, there was an increase of proteins related to cell adhesion, extracellular matrix organization, and membrane transport. Simultaneously, we detected a decrease in the number of proteins involved in oxidative phosphorylation, ion transport, and cytoskeleton organization (Figure 4e). The most surprising finding was the identification of increased APP levels in both human grafted cells and mouse host cells.

Considering these findings, we decided to compare the abundance of proteins involved in the AD-related KEGG pathway (hsa05010) across the four systems identified in our study, to map out the cellular origin, progression, and prion-like transmission of AD pathogenesis: ADHS, human cells in the ADHG, ADHPMBT and ADHG-MT (Figure 4f). Several proteins involved in lysosomal degradation, endocytosis, APP processing, oxidative phosphorylation, serine and glutathione metabolism, RNA processing and splicing were commonly dysregulated in the 4 systems. At the same time, proteins involved in endocytosis, serine metabolism, cytoskeleton reorganization, and oxidative phosphorylation were commonly dysregulated in human cells in the ADHG and ADHPMBT. In addition, the mitochondrial protein ETFDH involved in oxidative phosphorylation and the postsynaptic protein IQSEC2 involved in endocytosis, were commonly decreased in ADHS, human cells in the ADHG and ADHG-MT. Importantly, we identified APP as a protein commonly increased in human cells in ADHG, ADHPMBT and ADHG-MT (Figure 4f).

Altogether, the data suggest that different mechanisms are at play at different stages of AD pathology represented by the 4 systems, and that APP is central to AD cellular pathogenesis.

## Discussion

Most AD cases have an idiopathic origin. Transgenic AD animals have been the most common models to develop therapeutic strategies. Yet, they cannot clearly inform on the cellular origin and progression of AD patient brain dysfunction, even if they rely on the neuronal expression of human gene variants that lead to early or late onset AD. Because human iPSCs harbor the genetic background of the patient they are derived from, they may serve as a better alternative to understand AD cellular pathogenesis and develop treatments [33]. For example, work employing patient iPSCs has showed that haplotypes are important modulators of genetic risk factors for late-onset AD [34]. Recently, we and others successfully utilized AD patient iPSCs to generate 2D neuronal cell cultures and more complex free-floating organoid or spheroid structures, to uncover AD cellular phenotypes and test therapeutic strategies [5, 7, 34-37]. Altogether, these studies highlighted cell type specific dysfunctions and heterogeneity of patient cellular phenotypes. However, when grown in vitro, iPSC-derived brain cells lack the microenvironment that exists in the brain, preventing a more patient-like pathophysiological analysis of cellular phenotypes. To overcome this limitation several studies contemplated the possibility to study human cellular pathogenesis in humanized models. The human-rodent chimeric models allowed for the study of the effect of the AD brain microenvironment on human neurons [14] and glia [15].

Here, we thought to examine AD cell autonomous dysfunction in the healthy brain microenvironment, that naturally provides support to neurons and waste clearance, and to study the effect of the AD grafted brain cells on surrounding non-diseased host cells. We first demonstrated that the human grafted cells survived in the host mouse brain. Using LC-MS/MS, we observed increased abundance of APP and Tau, and the proteins involved in their processing, in ADHG compared to control. The proteomic changes were coupled to increased β-sheet structures and Aβ42 peptides. Yet, senile plaques measured by Congo red and amyloid antibody H31L21 staining were absent. We speculate that this could be attributed to the highly functional waste clearance system of the healthy host brain [38]. Another possibility lies in the fact that cell transplantation was performed in young mice aged 6-8 weeks. Najm and colleagues recently showed Aβ aggregation in iPSC-derived cortical neurons carrying ApoE4 risk factor, 7 months post-transplantation into the hippocampus of mice grafted when aged 6 months [13]. Alternatively, senile plaque formation may require longer period to form for the gene variant we studied. Indeed, plaques start to develop in the Thy1-APPLon mice when aged 10 months [39]. These points should be addressed in future studies and include the analysis of cellular phenotypes over longer periods, and with grafted cells having different genetic backgrounds.

Human astrocytes have emerged as a major player in AD [36, 40-43]. Not only these cells can become reactive when exposed to Aβ in vivo [15], and their secretion of cytokines enhanced following stimulation [36], their ability to uptake glutamate, release lactate and propagate calcium is altered if carrying a genetic variant leading to early onset AD [36]. We previously observed similar features when Parkinsonian and control iPSC-derived astrocytes were exposed to alpha-synuclein protein aggregates [44]. Here, ADHG contained fewer human astrocytes that CHG. However, ADHG contained many more host astrocytes. Possibly, the ADHG had attracted and activated a greater number of host astrocytes due to higher level of Aβ42 in the ADHG compared to control. Future work should explore whether cell autonomous dysfunctions exist in APP p.V717I astrocytes, in addition to the increase in β-sheet structure formation previously identified [7].

Here, the grafts were composed of both human and mouse cells. Thus, we had to conduct a database search of the LC-MS/MS spectra against the combined *Homo sapiens* and *Mus musculus* databases (Uniprot) to quantify species-specific unique peptides and investigate them individually. Proteomic analysis revealed profound cellular alterations in ADHG. Mitochondria homeostasis is one of the earliest impaired intracellular processes in AD [45–47]. In line with this, we identified several mitochondria associated proteins whose levels were changed in ADHG. Importantly, we observed an increase in protein kinases and proteases, including but not limited to GSK3B, GSK3A, BACE1, MARK1 and MARK3. These proteins, over-activity of which accounts for memory impairment, tau-hyperphosphorylation and increased Aβ production [48] increase with age and in AD [49–51]. Additionally, we observed an important decrease in several synaptic proteins in the ADHG, which is in line with the observations that synaptic pathology is an early feature of AD [52, 53].

Evidence suggests that both Aβ and Tau protein aggregates can spread throughout the brain in prion-like ways to trigger cellular dysfunction and seed de novo aggregation in recipient cells [30, 54-56]. In line with this, secretomes of AD iPSC-derived neurons delivered to the adult rat hippocampus resulted in impaired synaptic plasticity via a common pathway that is mediated by cellular prion proteins [32]. Interestingly, we identified several common proteomic alterations in both human and mouse proteomes, suggesting transfer of cellular pathology from transplanted human AD hippocampal cells to host mouse cells. One of the commonly dysregulated proteins was APP, accumulation of which occurs early in the cascade of events that results in senile plaque formation in AD brain. Interestingly, not only pathogenic variations in APP but also higher levels of wildtype APP either due to chromosomal translocation (trisomy 21) or APP gene locus duplication, lead to early onset AD [57–59]. We speculate that increased APP levels are induced by Aβ peptides secreted by the human AD cells, which is in line with earlier reports [60]. While future studies may elucidate the mechanism leading to increased APP level in host cells, current effort should aim at developing strategies to lower APP level which may be a relevant therapeutic avenue to slow down AD progression.

### Limitations of this study

We acknowledge several limitations in our study. Recent single cell/nuclei sequencing data clearly indicate that genes whose variants lead to early onset AD or genetic risk factors for late onset AD, are prominently expressed in glia in the human brain (https://portal.brain-map.org/atlases-and-data/rnaseq and https://www.brainrnaseq.org/). For instance, the expression of Presenilin 1 and APP genes is higher in oligodendrocytes than neurons. This suggest that oligodendrocyte dysfunction could play a role in early onset AD. White matter hyperintensities have been found elevated among individuals with early onset AD up to 20 years before the expected onset of symptoms [61]. Despite our capability to generate all brain cell types from human iPSCs [7, 44, 62], we have not yet developed a protocol to generate highly regionalized hippocampal spheroids containing neurons, astrocytes, and oligodendrocytes. Such a model system could allow for the examination of myelination defects and demyelination, both in vitro and in vivo. Likewise, microglia, cells of mesodermal/mesenchymal origin, are important in the formation of senile plaques [63]. TREM2 pathogenic variants supposedly lead to AD via decreased Aβ clearance due to defective phagocytosis by microglia [64]. The incorporation of iPSC-derived microglia could provide further information into neuronal phenotypes [65].

Additionally, some animals experienced epileptic seizures; for ethical reasons, these had to be put down, which is also the reason why we ended our study 6 months posttransplantation. We speculate seizures to be consequent to dysregulation of local hippocampal networks induced by the graft over time. In future studies, we could transplant either fewer brain cells or purified non-dividing cell types and follow how cellular pathology would develop in them over longer periods. Alternatively, we could graft whole brain spheroids by cranial window surgery, as performed for organoids [66, 67]. This would allow us to perform two-photon imaging studies to reveal neuronal network activity of the AD patient transplanted cells.

Finally, the hippocampal tissues and iPSC-derived brain cells analyzed originated from different patients. Whereas iPSC-derived brain cells carried the APP London mutation, the post-mortem brain tissue samples originated from idiopathic AD cases carrying ApoE3/E3. This is a major drawback since the progression of the AD cellular pathways from iPSC-derived models to patient postmortem tissue could only be predicted and not ascertained. Nevertheless, it could be argued that since all AD cellular pathways and network dysfunctions converge towards the same cellular hallmarks, the tissue analyzed represented the best available resource for the study. This issue could be circumvented if APP London mutation brains become available.

## Conclusions

This study highlights the promise of studying patient cellular pathogenesis in vivo, in experimental humanized rodent models. By transplanting iPSC-derived brain cells into the mouse brain, we were able to demonstrate what cellular pathways and networks are altered in young AD patient hippocampal cells and compare them to those present in ADHS grown in vitro and patient post-mortem tissue. Importantly, we showed a putative transfer of cellular pathology from diseased cells to healthy ones, like what has been proposed for other neurodegenerative diseases [68–70]. This new human/mouse chimeric model represents a powerful resource to better study the mechanisms underlying the cellular origins and progression of human AD pathogenesis, *in vivo*. We anticipate the future use of humanized rodent models to develop and test therapeutic strategies for AD, and importantly, validate target engagement prior to moving experimental therapies to clinical trials.

## List of abbreviations

Aβ: amyloid-β
AD: Alzheimer’s disease
ADHG: AD hippocampal grafts
ADHG-MT: mouse tissue adjacent to the AD hippocampal graft
ADHPMBT: AD hippocampal postmortem brain tissue
ADHS: AD hippocampal spheroids
ApoE4: Alipoprotein E
APP: amyloid precursor protein
BACE1: Beta-Secretase 1
CHG: control hippocampal grafts
Da: Dalton
DAPI: 4’,6-diamidino-2-phenylindole
FDR: false discovery rate
FGF2: Fibroblast growth factor 2
FTIR: Fourier transform infrared
GFAP: glial fibrillary acidic protein
GSK3A: glycogen synthase kinase-3 alpha
GSK3B: glycogen synthase kinase-3 beta
IBA1: ionized calcium binding adaptor molecule 1
iPSC: induced pluripotent stem cells
IT: injection time
KEGG: Kyoto Encyclopedia of Genes and Genomes
LC-MS/MS: liquid chromatography-tandem mass spectrometry
MAP2: Microtubule-associated protein 2
MARK1: Microtubule affinity regulating kinase 1
MARK3: Microtubule affinity regulating kinase 3
MSD: Meso Scale Discovery
NDS: normal donkey serum
PBS: Phosphate-buffered saline
PBST: Phosphate-buffered saline tween
PD: Proteome Discoverer
PET: Positron emission tomography
PFA: Paraformaldehyde
PVDF: Polyvinylidene fluoride
RAG-2: recombination activation gene 2
RT: room temperature
TBR1: T-Box Brain Transcription Factor 1
TEAB: Triethylammonium bicarbonate
TREM2: Triggering receptors expressed on myeloid cells 2
ZBTB20: Zinc Finger and BTB Domain Containing 20
18F: flutemetamol tracer

## Declarations

### Ethical Approval and Consent to participate

All experimental procedures were conducted in accordance with the European Union Directive (2010/63/EU) about animal rights and were approved by the committees for the use of laboratory animals at Lund University and the Swedish Board of Agriculture.

### Consent for publication

Not applicable.

### Availability of supporting data

The mass spectrometry data of ADHS have been deposited to the ProteomeXchange Consortium via the PRIDE partner repository with the dataset identifier PXD012524. The mass spectrometry data of ADGH and ADHPMBT are available from the corresponding author upon request. Omics data are also available as supplementary material.

### Authors’ contribution

Y.P., E.V. and L.R. conceived the experiments and wrote the manuscript; All authors performed or assisted with the experiments; and provided reagents, expertise and conducted a critical analysis the data and review of the manuscript. All authors read and approved the final manuscript.

## Acknowledgments

We are extremely thankful to Marianne Juhlin and other members of the CSC laboratory for their outstanding technical assistance. We are also thankful to the Netherlands Brain Bank for providing patient post-mortem tissue samples. Lund University Bioimaging Centre (LBIC), Lund University, is gratefully acknowledged for providing experimental resources.

## References

1. Blennow K, de Leon MJ, Zetterberg H: Alzheimer’s disease. Lancet 2006, 368:387–403.

2. Oakley H, Cole SL, Logan S, Maus E, Shao P, Craft J, Guillozet-Bongaarts A, Ohno M, Disterhoft J, Van Eldik L, et al: Intraneuronal beta-amyloid aggregates, neurodegeneration, and neuron loss in transgenic mice with five familial Alzheimer’s disease mutations: potential factors in amyloid plaque formation. J Neurosci 2006, 26:10129–10140.

3. Dimos JT, Rodolfa KT, Niakan KK, Weisenthal LM, Mitsumoto H, Chung W, Croft GF, Saphier G, Leibel R, Goland R, et al: Induced pluripotent stem cells generated from patients with ALS can be differentiated into motor neurons. Science 2008, 321:1218–1221.

4. Sasaguri H, Hashimoto S, Watamura N, Sato K, Takamura R, Nagata K, Tsubuki S, Ohshima T, Yoshiki A, Sato K, et al: Recent Advances in the Modeling of Alzheimer’s Disease. Front Neurosci 2022, 16:807473.

5. Konttinen H, Cabral-da-Silva MEC, Ohtonen S, Wojciechowski S, Shakirzyanova A, Caligola S, Giugno R, Ishchenko Y, Hernandez D, Fazaludeen MF, et al: PSEN1DeltaE9, APPswe, and APOE4 Confer Disparate Phenotypes in Human iPSC-Derived Microglia. Stem Cell Reports 2019, 13:669–683.

6. Kondo T, Asai M, Tsukita K, Kutoku Y, Ohsawa Y, Sunada Y, Imamura K, Egawa N, Yahata N, Okita K, et al: Modeling Alzheimer’s disease with iPSCs reveals stress phenotypes associated with intracellular Abeta and differential drug responsiveness. Cell Stem Cell 2013, 12:487–496.

7. Pomeshchik Y, Klementieva O, Gil J, Martinsson I, Hansen MG, de Vries T, Sancho-Balsells A, Russ K, Savchenko E, Collin A, et al: Human iPSC-Derived Hippocampal Spheroids: An Innovative Tool for Stratifying Alzheimer Disease Patient-Specific Cellular Phenotypes and Developing Therapies. Stem Cell Reports 2020, 15:256–273.

8. Penney J, Ralvenius WT, Tsai LH: Modeling Alzheimer’s disease with iPSC-derived brain cells. Mol Psychiatry 2020, 25:148–167.

9. Verheijen MCT, Krauskopf J, Caiment F, Nazaruk M, Wen QF, van Herwijnen MHM, Hauser DA, Gajjar M, Verfaillie C, Vermeiren Y, et al: iPSC-derived cortical neurons to study sporadic Alzheimer disease: A transcriptome comparison with post-mortem brain samples. Toxicol Lett 2022, 356:89–99.

10. Zhao J, Fu Y, Yamazaki Y, Ren Y, Davis MD, Liu CC, Lu W, Wang X, Chen K, Cherukuri Y, et al: APOE4 exacerbates synapse loss and neurodegeneration in Alzheimer’s disease patient iPSC-derived cerebral organoids. Nat Commun 2020, 11:5540.

11. Windrem MS, Osipovitch M, Liu Z, Bates J, Chandler-Militello D, Zou L, Munir J, Schanz S, McCoy K, Miller RH, et al: Human iPSC Glial Mouse Chimeras Reveal Glial Contributions to Schizophrenia. Cell Stem Cell 2017, 21:195–208 e196.

12. Osipovitch M, Asenjo Martinez A, Mariani JN, Cornwell A, Dhaliwal S, Zou L, Chandler-Militello D, Wang S, Li X, Benraiss SJ, et al: Human ESC-Derived Chimeric Mouse Models of Huntington’s Disease Reveal Cell-Intrinsic Defects in Glial Progenitor Cell Differentiation. Cell Stem Cell 2019, 24:107–122 e107.

13. Najm R, Zalocusky KA, Zilberter M, Yoon SY, Hao Y, Koutsodendris N, Nelson M, Rao A, Taubes A, Jones EA, Huang Y: In Vivo Chimeric Alzheimer’s Disease Modeling of Apolipoprotein E4 Toxicity in Human Neurons. Cell Rep 2020, 32:107962.

14. Espuny-Camacho I, Arranz AM, Fiers M, Snellinx A, Ando K, Munck S, Bonnefont J, Lambot L, Corthout N, Omodho L, et al: Hallmarks of Alzheimer’s Disease in Stem-Cell-Derived Human Neurons Transplanted into Mouse Brain. Neuron 2017, 93:1066–1081 e1068.

15. Preman P, Tcw J, Calafate S, Snellinx A, Alfonso-Triguero M, Corthout N, Munck S, Thal DR, Goate AM, De Strooper B, Arranz AM: Human iPSC-derived astrocytes transplanted into the mouse brain undergo morphological changes in response to amyloid-beta plaques. Mol Neurodegener 2021, 16:68.

16. Zhou Y, Zhou B, Pache L, Chang M, Khodabakhshi AH, Tanaseichuk O, Benner C, Chanda SK: Metascape provides a biologist-oriented resource for the analysis of systems-level datasets. Nat Commun 2019, 10:1523.

17. Braak H, Braak E: Neuropathological stageing of Alzheimer-related changes. Acta Neuropathol 1991, 82:239–259.

18. Matsuda H, Ito K, Ishii K, Shimosegawa E, Okazawa H, Mishina M, Mizumura S, Ishii K, Okita K, Shigemoto Y, et al: Quantitative Evaluation of (18)F-Flutemetamol PET in Patients With Cognitive Impairment and Suspected Alzheimer’s Disease: A Multicenter Study. Front Neurol 2020, 11:578753.

19. Bouter C, Bouter Y: (18)F-FDG-PET in Mouse Models of Alzheimer’s Disease. Front Med (Lausanne) 2019, 6:71.

20. Klementieva O, Willen K, Martinsson I, Israelsson B, Engdahl A, Cladera J, Uvdal P, Gouras GK: Pre-plaque conformational changes in Alzheimer’s disease-linked Abeta and APP. Nat Commun 2017, 8:14726.

21. Wu M, Zhang M, Yin X, Chen K, Hu Z, Zhou Q, Cao X, Chen Z, Liu D: The role of pathological tau in synaptic dysfunction in Alzheimer’s diseases. Transl Neurodegener 2021, 10:45.

22. Kayed R, Head E, Sarsoza F, Saing T, Cotman CW, Necula M, Margol L, Wu J, Breydo L, Thompson JL, et al: Fibril specific, conformation dependent antibodies recognize a generic epitope common to amyloid fibrils and fibrillar oligomers that is absent in prefibrillar oligomers. Mol Neurodegener 2007, 2:18.

23. Delacourte A: General and dramatic glial reaction in Alzheimer brains. Neurology 1990, 40:33–37.

24. Reichenbach N, Delekate A, Plescher M, Schmitt F, Krauss S, Blank N, Halle A, Petzold GC: Inhibition of Stat3-mediated astrogliosis ameliorates pathology in an Alzheimer’s disease model. EMBO Mol Med 2019, 11.

25. Toral-Rios D, Patino-Lopez G, Gomez-Lira G, Gutierrez R, Becerril-Perez F, Rosales-Cordova A, Leon-Contreras JC, Hernandez-Pando R, Leon-Rivera I, Soto-Cruz I, et al: Activation of STAT3 Regulates Reactive Astrogliosis and Neuronal Death Induced by AbetaO Neurotoxicity. Int J Mol Sci 2020, 21.

26. Kwart D, Gregg A, Scheckel C, Murphy EA, Paquet D, Duffield M, Fak J, Olsen O, Darnell RB, Tessier-Lavigne M: A Large Panel of Isogenic APP and PSEN1 Mutant Human iPSC Neurons Reveals Shared Endosomal Abnormalities Mediated by APP beta-CTFs, Not Abeta. Neuron 2019, 104:256–270 e255.

27. Bastin G, Heximer SP: Rab family proteins regulate the endosomal trafficking and function of RGS4. J Biol Chem 2013, 288:21836–21849.

28. Tam SY, Lilla JN, Chen CC, Kalesnikoff J, Tsai M: RabGEF1/Rabex-5 Regulates TrkA-Mediated Neurite Outgrowth and NMDA-Induced Signaling Activation in NGF-Differentiated PC12 Cells. PLoS One 2015, 10:e0142935.

29. Nagano M, Toshima JY, Siekhaus DE, Toshima J: Rab5-mediated endosome formation is regulated at the trans-Golgi network. Commun Biol 2019, 2:419.

30. Aoyagi A, Condello C, Stohr J, Yue W, Rivera BM, Lee JC, Woerman AL, Halliday G, van Duinen S, Ingelsson M, et al: Abeta and tau prion-like activities decline with longevity in the Alzheimer’s disease human brain. Sci Transl Med 2019, 11.

31. Gomez-Gutierrez R, Morales R: The prion-like phenomenon in Alzheimer’s disease: Evidence of pathology transmission in humans. PLoS Pathog 2020, 16:e1009004.

32. Hu NW, Corbett GT, Moore S, Klyubin I, O’Malley TT, Walsh DM, Livesey FJ, Rowan MJ: Extracellular Forms of Abeta and Tau from iPSC Models of Alzheimer’s Disease Disrupt Synaptic Plasticity. Cell Rep 2018, 23:1932–1938.

33. Okano H, Morimoto S: iPSC-based disease modeling and drug discovery in cardinal neurodegenerative disorders. Cell Stem Cell 2022, 29:189–208.

34. Tcw J, Qian L, Pipalia NH, Chao MJ, Liang SA, Shi Y, Jain BR, Bertelsen SE, Kapoor M, Marcora E, et al: Cholesterol and matrisome pathways dysregulated in astrocytes and microglia. Cell 2022, 185:2213–2233 e2225.

35. Muratore CR, Rice HC, Srikanth P, Callahan DG, Shin T, Benjamin LN, Walsh DM, Selkoe DJ, Young-Pearse TL: The familial Alzheimer’s disease APPV717I mutation alters APP processing and Tau expression in iPSC-derived neurons. Hum Mol Genet 2014, 23:3523–3536.

36. Oksanen M, Petersen AJ, Naumenko N, Puttonen K, Lehtonen S, Gubert Olive M, Shakirzyanova A, Leskela S, Sarajarvi T, Viitanen M, et al: PSEN1 Mutant iPSC-Derived Model Reveals Severe Astrocyte Pathology in Alzheimer’s Disease. Stem Cell Reports 2017, 9:1885–1897.

37. Shimada H, Sato Y, Sasaki T, Shimozawa A, Imaizumi K, Shindo T, Miyao S, Kiyama K, Kondo T, Shibata S, et al: A next-generation iPSC-derived forebrain organoid model of tauopathy with tau fibrils by AAV-mediated gene transfer. Cell Rep Methods 2022, 2:100289.

38. Nedergaard M, Goldman SA: Glymphatic failure as a final common pathway to dementia. Science 2020, 370:50–56.

39. Van Dorpe J, Smeijers L, Dewachter I, Nuyens D, Spittaels K, Van Den Haute C, Mercken M, Moechars D, Laenen I, Kuiperi C, et al: Prominent cerebral amyloid angiopathy in transgenic mice overexpressing the london mutant of human APP in neurons. Am J Pathol 2000, 157:1283–1298.

40. Albert K, Niskanen J, Kalvala S, Lehtonen S: Utilising Induced Pluripotent Stem Cells in Neurodegenerative Disease Research: Focus on Glia. Int J Mol Sci 2021, 22.

41. Jones VC, Atkinson-Dell R, Verkhratsky A, Mohamet L: Aberrant iPSC-derived human astrocytes in Alzheimer’s disease. Cell Death Dis 2017, 8:e2696.

42. Habib N, McCabe C, Medina S, Varshavsky M, Kitsberg D, Dvir-Szternfeld R, Green G, Dionne D, Nguyen L, Marshall JL, et al: Disease-associated astrocytes in Alzheimer’s disease and aging. Nat Neurosci 2020, 23:701–706.

43. Salcedo C, Andersen JV, Vinten KT, Pinborg LH, Waagepetersen HS, Freude KK, Aldana BI: Functional Metabolic Mapping Reveals Highly Active Branched-Chain Amino Acid Metabolism in Human Astrocytes, Which Is Impaired in iPSC-Derived Astrocytes in Alzheimer’s Disease. Front Aging Neurosci 2021, 13:736580.

44. Russ K, Teku G, Bousset L, Redeker V, Piel S, Savchenko E, Pomeshchik Y, Savistchenko J, Stummann TC, Azevedo C, et al: TNF-alpha and alpha-synuclein fibrils differently regulate human astrocyte immune reactivity and impair mitochondrial respiration. Cell Rep 2021, 34:108895.

45. Wang W, Zhao F, Ma X, Perry G, Zhu X: Mitochondria dysfunction in the pathogenesis of Alzheimer’s disease: recent advances. Mol Neurodegener 2020, 15:30.

46. Cenini G, Voos W: Mitochondria as Potential Targets in Alzheimer Disease Therapy: An Update. Front Pharmacol 2019, 10:902.

47. Kobro-Flatmoen A, Lagartos-Donate MJ, Aman Y, Edison P, Witter MP, Fang EF: Re-emphasizing early Alzheimer’s disease pathology starting in select entorhinal neurons, with a special focus on mitophagy. Ageing Res Rev 2021, 67:101307.

48. Hooper C, Killick R, Lovestone S: The GSK3 hypothesis of Alzheimer’s disease. J Neurochem 2008, 104:1433–1439.

49. Ma T: GSK3 in Alzheimer’s disease: mind the isoforms. J Alzheimers Dis 2014, 39:707–710.

50. Lauretti E, Dincer O, Pratico D: Glycogen synthase kinase-3 signaling in Alzheimer’s disease. Biochim Biophys Acta Mol Cell Res 2020, 1867:118664.

51. Chudobova J, Zempel H: Microtubule affinity regulating kinase (MARK/Par1) isoforms differentially regulate Alzheimer-like TAU missorting and Abeta-mediated synapse pathology. Neural Regen Res 2023, 18:335–336.

52. Forner S, Baglietto-Vargas D, Martini AC, Trujillo-Estrada L, LaFerla FM: Synaptic Impairment in Alzheimer’s Disease: A Dysregulated Symphony. Trends Neurosci 2017, 40:347–357.

53. Scheff SW, Price DA, Schmitt FA, Mufson EJ: Hippocampal synaptic loss in early Alzheimer’s disease and mild cognitive impairment. Neurobiol Aging 2006, 27:1372–1384.

54. Ayers JI, Giasson BI, Borchelt DR: Prion-like Spreading in Tauopathies. Biol Psychiatry 2018, 83:337–346.

55. Condello C, Stoehr J: Abeta propagation and strains: Implications for the phenotypic diversity in Alzheimer’s disease. Neurobiol Dis 2018, 109:191–200.

56. Roos TT, Garcia MG, Martinsson I, Mabrouk R, Israelsson B, Deierborg T, Kobro-Flatmoen A, Tanila H, Gouras GK: Neuronal spreading and plaque induction of intracellular Abeta and its disruption of Abeta homeostasis. Acta Neuropathol 2021, 142:669–687.

57. Sleegers K, Brouwers N, Gijselinck I, Theuns J, Goossens D, Wauters J, Del-Favero J, Cruts M, van Duijn CM, Van Broeckhoven C: APP duplication is sufficient to cause early onset Alzheimer’s dementia with cerebral amyloid angiopathy. Brain 2006, 129:2977–2983.

58. Fortea J, Zaman SH, Hartley S, Rafii MS, Head E, Carmona-Iragui M: Alzheimer’s disease associated with Down syndrome: a genetic form of dementia. Lancet Neurol 2021, 20:930–942.

59. Rovelet-Lecrux A, Hannequin D, Raux G, Le Meur N, Laquerriere A, Vital A, Dumanchin C, Feuillette S, Brice A, Vercelletto M, et al: APP locus duplication causes autosomal dominant early-onset Alzheimer disease with cerebral amyloid angiopathy. Nat Genet 2006, 38:24–26.

60. Davis-Salinas J, Van Nostrand WE: Amyloid beta-protein aggregation nullifies its pathologic properties in cultured cerebrovascular smooth muscle cells. J Biol Chem 1995, 270:20887–20890.

61. Nasrabady SE, Rizvi B, Goldman JE, Brickman AM: White matter changes in Alzheimer’s disease: a focus on myelin and oligodendrocytes. Acta Neuropathol Commun 2018, 6:22.

62. Azevedo C, Teku G, Pomeshchik Y, Reyes JF, Chumarina M, Russ K, Savchenko E, Hammarberg A, Lamas NJ, Collin A, et al: Parkinson’s disease and multiple system atrophy patient iPSC-derived oligodendrocytes exhibit alpha-synuclein-induced changes in maturation and immune reactive properties. Proc Natl Acad Sci U S A 2022, 119:e2111405119.

63. Venegas C, Kumar S, Franklin BS, Dierkes T, Brinkschulte R, Tejera D, Vieira-Saecker A, Schwartz S, Santarelli F, Kummer MP, et al: Microglia-derived ASC specks crossseed amyloid-beta in Alzheimer’s disease. Nature 2017, 552:355–361.

64. Gratuze M, Leyns CEG, Holtzman DM: New insights into the role of TREM2 in Alzheimer’s disease. Mol Neurodegener 2018, 13:66.

65. Fagerlund I, Dougalis A, Shakirzyanova A, Gomez-Budia M, Pelkonen A, Konttinen H, Ohtonen S, Fazaludeen MF, Koskuvi M, Kuusisto J, et al: Microglia-like Cells Promote Neuronal Functions in Cerebral Organoids. Cells 2021, 11.

66. Mansour AA, Goncalves JT, Bloyd CW, Li H, Fernandes S, Quang D, Johnston S, Parylak SL, Jin X, Gage FH: An in vivo model of functional and vascularized human brain organoids. Nat Biotechnol 2018, 36:432–441.

67. Revah O, Gore F, Kelley KW, Andersen J, Sakai N, Chen X, Li MY, Birey F, Yang X, Saw NL, et al: Maturation and circuit integration of transplanted human cortical organoids. Nature 2022, 610:319–326.

68. Brundin P, Melki R: Prying into the Prion Hypothesis for Parkinson’s Disease. J Neurosci 2017, 37:9808–9818.

69. Gosset P, Camu W, Raoul C, Mezghrani A: Prionoids in amyotrophic lateral sclerosis. Brain Commun 2022, 4:fcac145.

70. Donnelly KM, Coleman CM, Fuller ML, Reed VL, Smerina D, Tomlinson DS, Pearce MMP: Hunting for the cause: Evidence for prion-like mechanisms in Huntington’s disease. Front Neurosci 2022, 16:946822.

