## Supplementary figures and images for "Proteomic analysis across patient iPSC-based models and human post-mortem hippocampal tissue reveals early cellular dysfunction, progression, and prion-like spread of Alzheimer’s disease pathogenesis"

### Figure S1

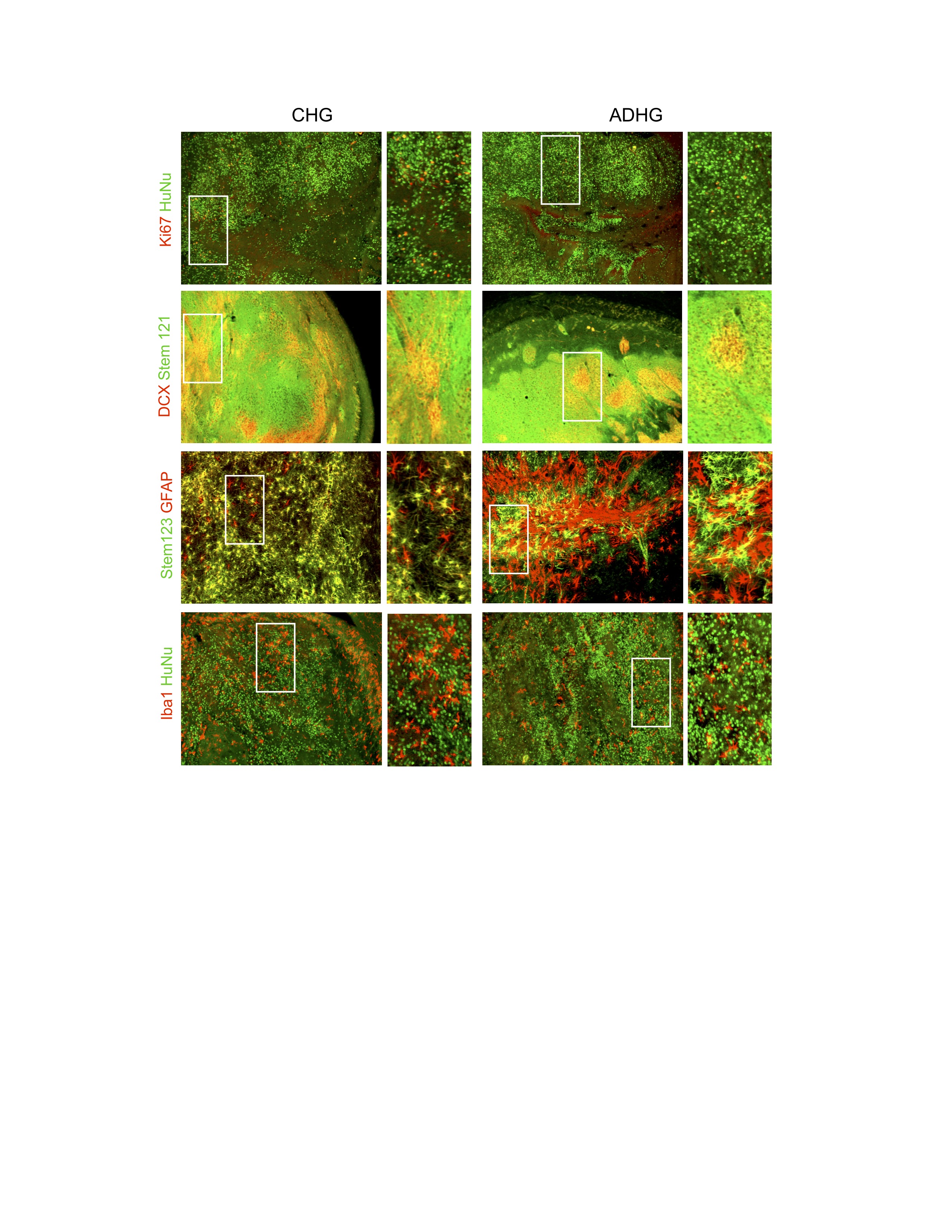
