## Supplementary material for "Proteomic analysis across patient iPSC-based models and human post-mortem hippocampal tissue reveals early cellular dysfunction, progression, and prion-like spread of Alzheimer’s disease pathogenesis": Table S6

**Supplementary Table 6. List of human post-mortem brain samples**

| **Gender** | **Age** | **Diagnosis** | **Braak stage** | **Lewy Bodies** | **Postmortem delay** |
| --- | --- | --- | --- | --- | --- |
| F | 69 | Alzheimer’s Disease | 5 C |  | 5:45 |
| F | 70 | Alzheimer’s Disease | 6 C |  | 4:30 |
| M | 56 | Alzheimer’s Disease | 6 C | 2 | 4:40 |
| M | 69 | Alzheimer’s Disease | 5 B |  | 7:10 |
| F | 85 | Control with cerebrovascular accident | 2 B |  | 7:50 |
| M | 93 | Non-demented control | 0 A |  | 7:40 |
| F | 75 | Non-demented control | 1 A |  | 9:10 |
