## Supplementary material for "Proteomic analysis across patient iPSC-based models and human post-mortem hippocampal tissue reveals early cellular dysfunction, progression, and prion-like spread of Alzheimer’s disease pathogenesis": Table S7

**Supplementary Table 7. Primary and secondary antibodies used for immunocyto- and immunohistochemistry.**

| **Antibody** | **Species** | **Type** | **Dilution** | **Manufacturer** | **Reference** |
| --- | --- | --- | --- | --- | --- |
| **Primary antibodies** | | | | |  |
| AT8 | Mouse | Monoclonal | 1:500 | Thermo Fisher | MN1020 |
| Doublecortin | Rabbit | Monoclonal | 1:400 | Cell Signalling | 4604 |
| GFAP | Rabbit | Polyclonal | 1:2000 | DAKO | Z0334 |
| H31L21 | Rabbit | Monoclonal | 1:500 | Thermo Fisher | 700254 |
| Human nuclear antigen (hNuclei) | Mouse | Monoclonal | 1:250 | Merck Millipore | MAB1281 |
| IBA1 | Rabbit | Polyclonal | 1:500 | WAKO | 019-19741 |
| KI67 | Rabbit | Polyclonal | 1:1000 | Thermo Fisher | PA5-19462 |
| MAP2 | Chicken | Polyclonal | 1:1000 | Abcam | AB92434 |
| PROX1 | Rabbit | Monoclonal | 1:400 | Abcam | ab199359 |
| STEM 121 | Mouse | Monoclonal | 1:500 | Takara Bio Inc. | Y40410 |
| STEM 123 | Mouse | Monoclonal | 1:500 | Takara Bio Inc. | Y40420 |
| TBR1 | Rabbit | Polyclonal | 1:500 | Merck Millipore | AB10554 |
| ZBTB20 | Rabbit | Polyclonal | 1:200 | Sigma Aldrich | HPA016815 |
| **Secondary antibodies** | | | |  |  |
| Anti-mouse Alexa Fluor® 488 | Donkey | Polyclonal | 1:400 | Thermo Fisher | A21202 |
| Anti-rabbit Alexa Fluor® 488 | Donkey | Polyclonal | 1:400 | Thermo Fisher | A21206 |
| Anti-mouse Alexa Fluor® 555 | Donkey | Polyclonal | 1:400 | Thermo Fisher | A31570 |
| Anti-rabbit Alexa Fluor® 555 | Donkey | Polyclonal | 1:400 | Thermo Fisher | A31572 |
| Anti-chicken Alexa Fluor® 647 | Donkey | Polyclonal | 1:400 | Merck Millipore | AP194SA6 |
