## Supplementary material for "Proteomic analysis across patient iPSC-based models and human post-mortem hippocampal tissue reveals early cellular dysfunction, progression, and prion-like spread of Alzheimer’s disease pathogenesis": Table S8

**Supplementary Table 8. Primary and secondary antibodies used for Western Blotting**

| **Antibody** | **Species** | **Type** | **Dilution** | **Manufacturer** | **Reference** |
| --- | --- | --- | --- | --- | --- |
| **Primary antibodies** | | | | |  |
| Actin | Mouse | Monoclonal | 1:1000 | Sigma | A5316 |
| AT8 | Mouse | Monoclonal | 1:1000 | Thermo Fisher | MN1020 |
| **Secondary antibodies** | | | |  |  |
| Anti-mouse IGG HRP-conjugated | Goat | Polyclonal | 1:1000 | R&D Systems | HAF007 |
